# Spatial neglect is causally linked to an attention-related area in macaque temporal cortex

**DOI:** 10.1101/410233

**Authors:** Amarender R. Bogadhi, Anil Bollimunta, David A. Leopold, Richard J. Krauzlis

**Affiliations:** Laboratory of Sensorimotor Research, National Eye Institute, National Institutes of Health; Laboratory of Neuropsychology, National Institute of Mental Health, National Institutes of Health; Neurophysiology Imaging Facility, National Institute of Mental Health, National Institute of Neurological Disorders and Stroke, National Eye Institute, National Institutes of Health

**Keywords:** spatial neglect, selective attention, superior colliculus, temporal cortex, fMRI, reversible inactivation, macaque

## Abstract

Spatial neglect is a common clinical syndrome involving disruption of the brain’s attention-related circuitry, including the dorsocaudal temporal cortex. While neglect is readily modeled in macaques, the absence of evidence for temporal cortex involvement has suggested a fundamental difference from humans. To map the neurological expression of spatial neglect in macaques, we measured attention-related fMRI activity across the cerebral cortex during experimental induction of neglect through reversible inactivation of the superior colliculus. During inactivation, monkeys exhibited hallmark attentional deficits of neglect in tasks using either moving or static stimuli. The behavioral deficits were accompanied by marked reductions in fMRI attentional modulation that were strongest in a small region on the floor of the superior temporal sulcus. Notably, direct inactivation of this mid-STS cortical region caused similar neglect-like spatial attention deficits. These results identify a putative macaque homolog to temporal cortex structures known to play a central role in human neglect.

**Highlights:**

- During neglect the mid-STS cortex shows largest reduction in attentional modulation
- The mid-STS cortex is selectively affected with static as well as dynamic stimuli
- Mid-STS cortex is affected during neglect caused by SC or FEF inactivation
- Inactivation of mid-STS cortex itself causes neglect without eye movement deficits

**In Brief:** Using fMRI in macaques performing attention tasks combined with reversible inactivations, Bogadhi et al. provide causal evidence that a region on the floor of the superior temporal sulcus is a crucial node in the cortical control of covert selective attention and a putative homolog to temporal cortex regions in humans implicated in spatial neglect.

## Introduction

In humans, spatial neglect is a common consequence of stroke damage, and is characterized by failures to orient, detect or respond to stimuli in the visual field opposite to the brain lesion in the absence of sensory processing or motor deficits [1-8].

The anatomy of spatial neglect is complex and includes multiple brain systems related to selective attention. Spatial neglect is often associated with damage to the inferior parietal and frontal cortices [9-13], although damage to subcortical structures in the telencephalon (e.g. striatum), thalamus (e.g. pulvinar) and midbrain (e.g. superior colliculus, SC) can lead to similar symptoms [14-16].

Recent work in humans has placed emphasis on the temporal cortex, with meta-analysis suggesting that its contribution to spatial neglect has been previously under-appreciated [17]. In particular, the dorsocaudal temporal cortex regions of the superior temporal gyrus (STG) and temporo-parietal junction (TPJ) have been hypothesized to play a particularly important role in spatial neglect [18-24]. These temporal cortical regions are also part of the functional network with frontal and parietal areas implicated in spatial attention in humans [25].

Understanding the brain mechanisms underlying spatial neglect requires systematic testing of hypotheses in animal models such as the macaque, whose perceptual specializations are similar to those in humans [26,27]. Attention-related deficits associated with visual neglect in humans have been demonstrated in the macaque following reversible inactivation of several homologous brain structures, such as the superior colliculus, pulvinar, parietal cortex (lateral intraparietal area, LIP) and prefrontal cortex (frontal eye fields, FEF) [28-32]. Attention-related areas have also been identified in the superior temporal sulcus of monkeys using fMRI [33-35]. However, direct comparison of fMRI results between humans and macaques suggests that monkeys might simply lack a homologue of the temporo-parietal junction [36], raising the possibility that the temporal cortical contribution to spatial neglect might be a specialization of the human cerebral cortex, and cannot be studied in macaques.

To determine brain regions functionally associated with spatial neglect in monkeys, we performed fMRI before and during midbrain-induced neglect in two monkeys performing visual selective attention tasks. We chose reversible inactivation of midbrain SC to induce spatial neglect because inactivation of this structure leads to robust attention-related deficits, which are a hallmark of neglect [29,37]. The broad coverage of fMRI provides a means to compare attention-related modulation throughout the brain under normal behavior (“control”) and following SC inactivation (“neglect”) to identify brain regions functionally affected during the induced neglect phenomenon.

We demonstrate that, across the monkey brain, the strongest unilateral suppression of attention-related modulation during midbrain-induced neglect was found in a circumscribed region of the mid-STS cortex. The reduction of attention-related modulation in this region during neglect did not depend on the stimulus feature used in the attention tasks, and the same temporal region was also affected by inactivation of prefrontal cortex. Moreover, in a direct causal test, we show that focal reversible inactivation of neuronal activity in this region of temporal cortex itself produced spatial neglect. These findings identify for the first time a region in the mid-STS cortex of macaque that is causally linked to hemi-spatial neglect and is functionally connected with both midbrain and prefrontal circuits for selective attention.

## Results

Two adult macaque monkeys were scanned using BOLD fMRI across repeated sessions while they performed a set of attention tasks previously demonstrated to identify attention-related modulation in humans and monkeys [35,38]. In each task, the color of the central cue indicated which stimulus should be attended covertly – the monkeys maintained strict central fixation and reported the relevant changes detected by releasing a joystick (Figure 1A-D). In the Baseline task, monkeys monitored the fixation stimulus to detect a possible luminance change. The Ignore task was similar but included peripheral visual stimuli that could change motion direction but were entirely task irrelevant. In the Attend task, monkeys monitored the same peripheral visual stimuli to detect possible changes in motion direction. All three tasks were matched in terms of overall difficulty; the hit rates for detecting relevant changes were comparable between Attend (Monkey #1: 69.4%; Monkey #2: 72.9%) and Ignore tasks (Monkey #1: 74.3%; Monkey #2: 65.1%). Importantly, previous fMRI studies in monkeys have shown that this central-vs-peripheral task design [35] activates peripheral visual field voxels in the same set of attention-related cortical areas as found with other task designs, such as left-vs-right cueing and visual search [33,34,39], indicating that identification of attention-related voxels in monkeys is highly reproducible across different labs and paradigms.

**Figure 1.**
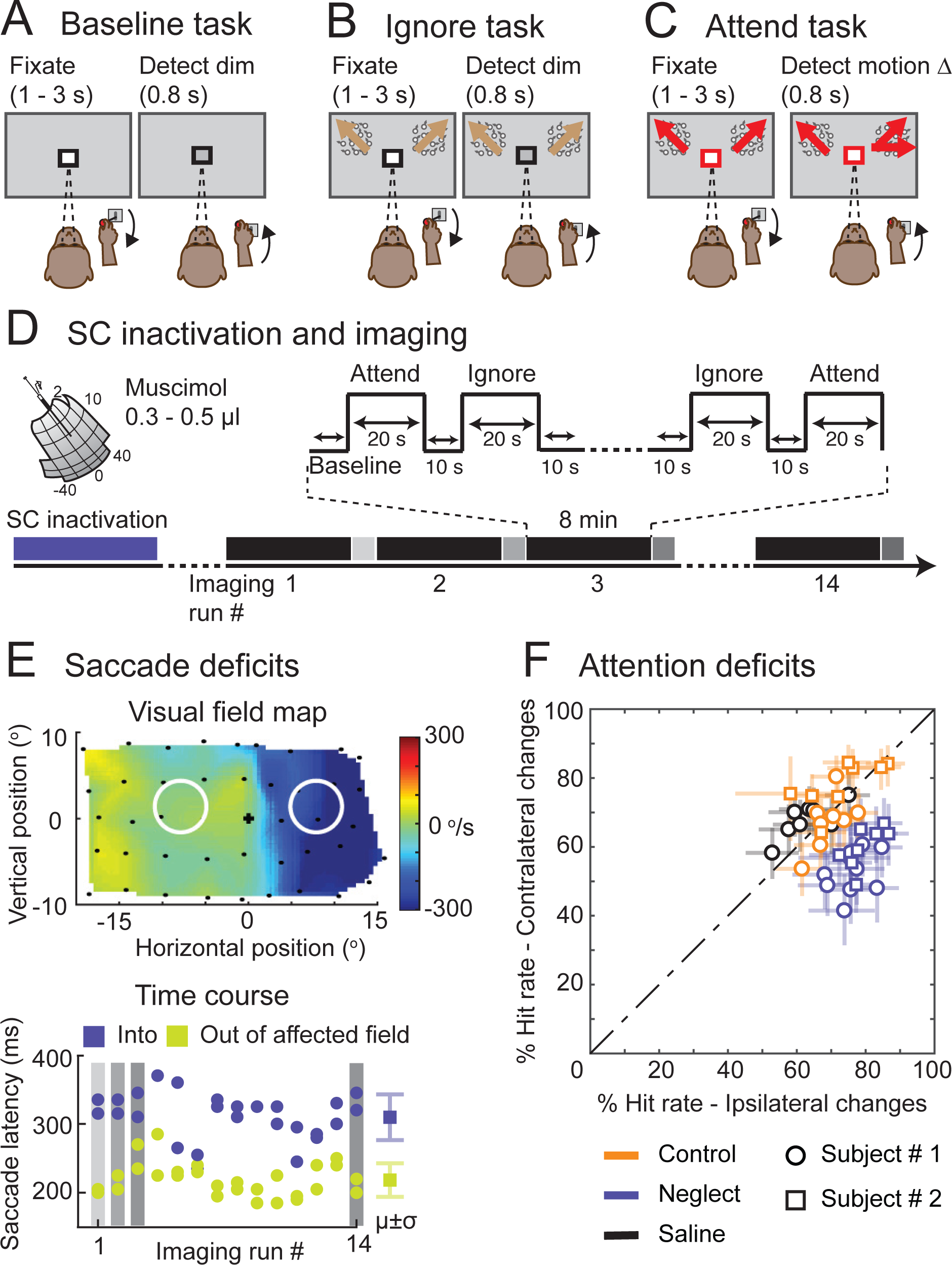
Behavioral tasks and midbrain-induced neglect (SC inactivation) (**A**) In the Baseline task, a black square around the central fixation spot instructed the monkey to monitor the fixation stimulus and detect luminance-change during a delay period of 1 – 3 seconds. (**B**) In the Ignore task, a black square around the central fixation spot instructed the monkey to monitor the fixation stimulus in the presence of peripheral motion distractors and detect luminance-change during a delay period of 1 – 3 seconds. Independent of the fixation spot dimming, one of the motion stimuli changed direction which the monkey ignored as part of the task. (**C**) In the Attend task, a red square around the central fixation spot instructed the monkey to monitor both the peripheral motion stimuli and detect a possible motion direction-change that occurred in one of the motion stimuli during a delay period of 1 – 3 seconds. The central fixation spot did not dim during Attend condition. In all tasks, the monkey reported the relevant change by releasing the joystick within 0.8 s for a juice reward. (**D**) Timeline of an example SC inactivation session and imaging. The session began with reversible inactivation of SC outside the scanner. In the scanner, the monkey performed all three tasks presented in a block design during an imaging run that lasted 8 minutes (black bars). At the end of each imaging run, the monkey performed visual guided saccades to targets in the affected and unaffected visual fields (grey bars). An average of 14 imaging runs were collected in an inactivation session. (**E**) The visual field map shows the effect of SC inactivation on the peak velocity of the saccades to targets in the visually guided saccade task (top panel). The blue region indicates the affected part of the visual field. Black dots indicate the retinotopic position of the saccade targets. The white circles indicate the positions of the motion stimuli in the Ignore and Attend tasks. The time course of muscimol effect was measured in the scanner using visual guided saccade task at the end of each imaging run (bottom panel). Blue dots indicate the latencies of saccades to targets in the affected visual field and green dots indicate the latencies of saccades to targets in the unaffected visual field. μ and σ indicate mean and standard deviation respectively. (**F**) Performance in Attend task is shown for control (green), neglect (SC inactivation; blue) and saline (black) sessions in both monkeys. % Hit rate for motion changes in the affected visual field (y-axis) is plotted against % hit rate for motion changes in the unaffected visual field (x-axis). Each data point represents a single session and error bars indicate 95% CI.

### Visual neglect induced by inactivation of superior colliculus

Visual neglect was induced by inactivation of the midbrain SC using microinjection of muscimol (0.3 – 0.5 μl into intermediate layers, left SC in monkey #1 and right SC in monkey #2, 18/42 fMRI sessions); in other fMRI sessions there was either no injection (16/42) or injection of saline (8/42) as a control. Overall, we obtained 4648 blocks of trials for each of the attention tasks (1792 with muscimol, 1928 with no injection, 928 with saline).

Consistent with previous findings, SC inactivation caused neglect-like deficits in the Attend task for contralateral stimulus changes [29,37]. During SC inactivation (Figure 1F), all but one session showed a significant reduction in hit rate for motion-direction changes in the affected contralateral visual field compared to the unaffected ipsilateral visual field (Chi-square proportion test; p<0.05). There was no difference in hit rates between the contralateral and ipsilateral visual fields (Chi-square proportion test; p>0.05) for the no-injection and saline sessions. Because the deficits caused by SC inactivation mimic the failures to detect contralateral targets observed in hemi-spatial neglect [25,40] we refer to these deficits as midbrain-induced neglect.

In addition to causing neglect, SC inactivation also affected the metrics of saccadic eye movements made to peripheral targets in a visually guided saccade task, and these changes in saccades provided a useful and independent measure to gauge the efficacy of the SC inactivation at multiple time points in each experiment. First, prior to each fMRI session, we mapped the spatial extent of the saccade deficits and confirmed that the portion of the visual field affected by SC inactivation fully overlapped the peripheral stimulus in the attention tasks but did not include the central visual field (Figure 1E). Second, we checked the effectiveness of muscimol at the peripheral stimulus location at the end of each imaging run (every 8 minutes) by having the monkeys perform a pair of visually guided saccade into and out of the affected visual field location (Figure 1E, bottom); for all data reported here, the latencies of saccades to targets into the affected field were significantly longer than the latencies of saccades to targets in the unaffected visual field (Wilcoxon rank sum test; p<0.05). In addition, during the fMRI sessions, we measured eye movements during the attention tasks to exclude the possibility that changes in the stability of fixation contributed to our imaging results.

### Mid-STS cortex showed largest reduction of attentional modulation during neglect

We first performed a voxel-wise analysis contrasting the BOLD measurements between the Attend and Ignore tasks, and projected the t-scores (Bonferroni corrected; p<0.05) onto partially inflated cortical surfaces for the hemisphere ipsilateral to the pending SC inactivation. During control sessions, there was significant modulation in areas previously implicated in visual attention [33-35], including visual areas, parts of the superior temporal sulcus (STS), and areas in parietal and frontal cortex (Figure 2A,C). During midbrain-induced neglect, several of these same areas showed reduced attention-related modulation, but the largest reductions were in portions of the mid-STS (Figure 2B,D). These effects were specific to the hemisphere associated with visual neglect; there was no evidence of suppressed attention-related modulation in the hemisphere opposite to the SC inactivation (Figure S1).

**Figure 2.**
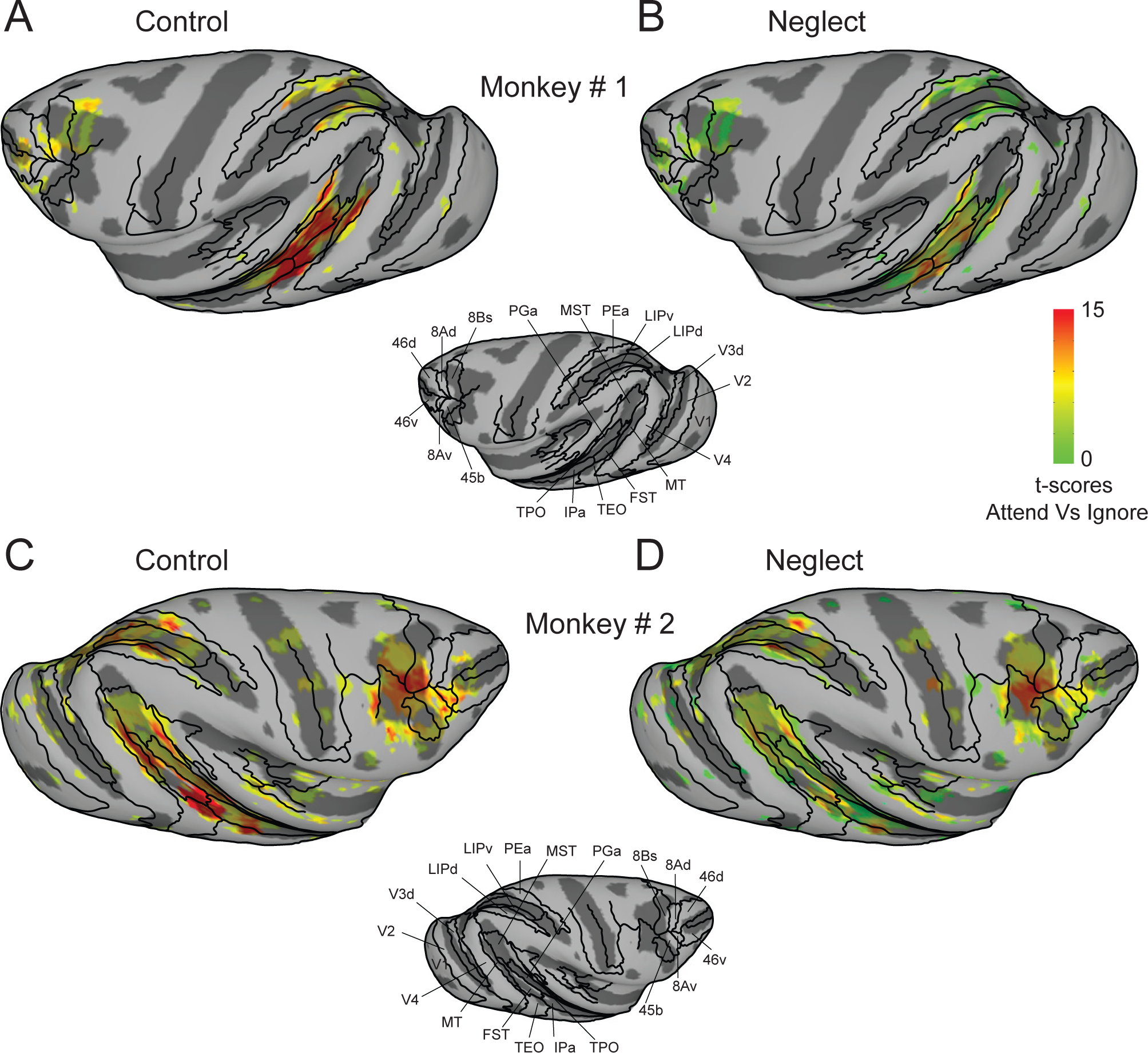
Cortical maps of attention-related modulation in control and neglect (SC inactivation) sessions. T-scores contrasting Attend and Ignore tasks were projected onto inflated cortical surfaces of D99 in each monkey’s native space along with anatomical boundaries (black contours). (**A,B**) Inflated cortical maps of t-scores show attention-related modulation in ipsilateral (left) hemisphere of monkey # 1 during control (**A**) and neglect (**B**). The maps were thresholded during control sessions (**A**) and the modulation for the same voxels is shown during the neglect sessions (**B**). (**C,D**) Same conventions as (**A,B**) but for monkey # 2. See also Figure S1.

To directly visualize the changes in attention-related modulation during neglect, we performed a voxel-wise ANOVA with two factors and two levels each (attend versus ignore, control versus neglect), similar to the analysis used in a previous human fMRI study of spatial attention deficits in neglect [25]. We then overlaid the interaction term F-statistic from the voxel-wise ANOVA onto partially inflated cortical surfaces of the ipsilateral hemispheres in both monkeys (Figure 3A,B), with negative values (blue) indicating decreases in modulation during neglect. Control analyses confirm that the thresholded maps in Figure 3 do not omit any regions whose modulation was reduced during neglect (Figure S2).

**Figure 3.**
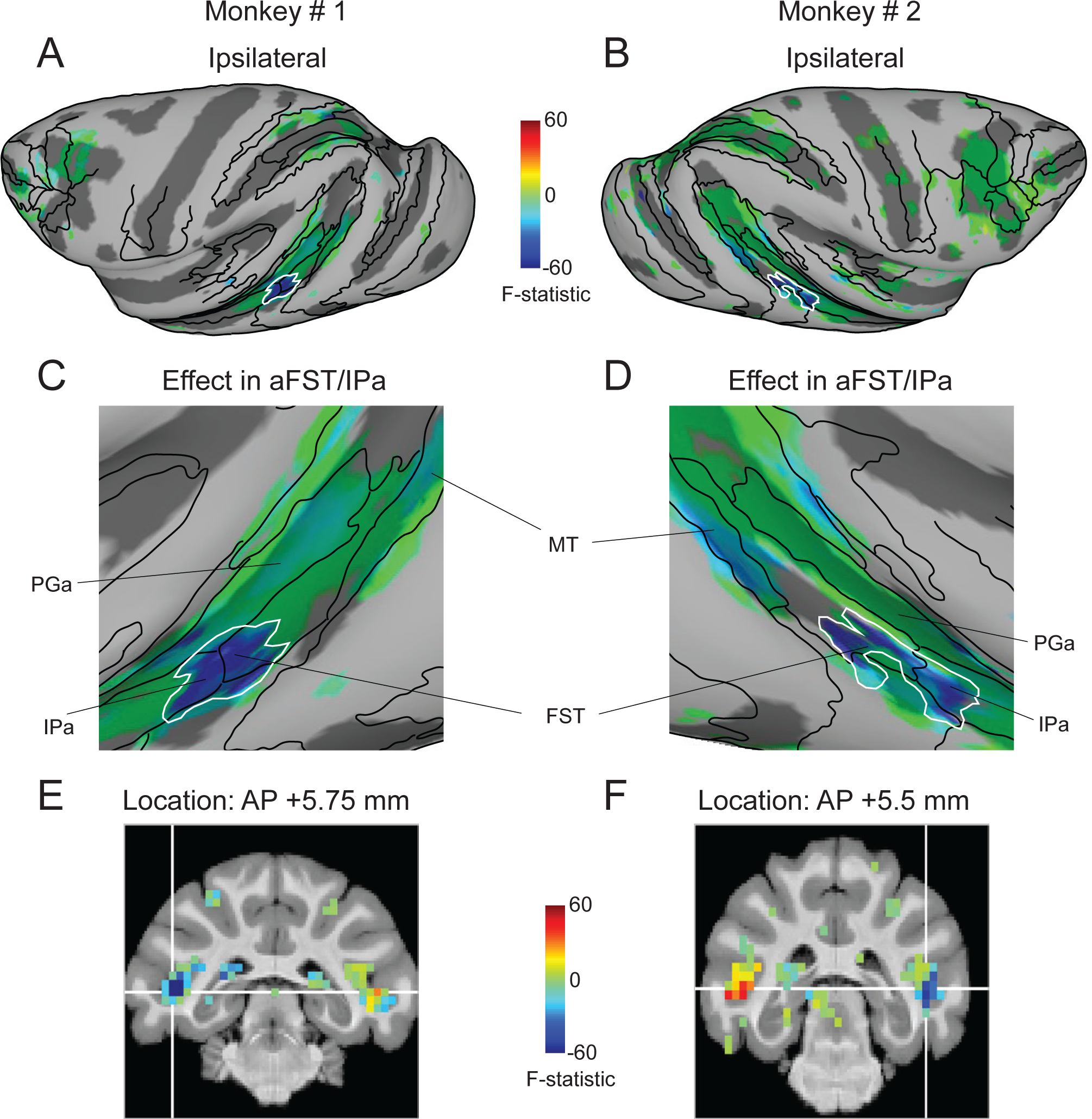
Midbrain-induced neglect linked to mid-STS cortex. (**A,B**) Inflated cortical maps of interaction term F-statistic show voxels whose modulation was affected during neglect in the ipsilateral hemisphere of monkey # 1 (**A**) and monkey # 2 (**B**). The effect during neglect is shown for the same voxels that showed significant modulation during control sessions (Figure 2A,C). The blue color indicates voxels whose modulation was reduced the most during neglect and the green colors indicate voxels with no reduction in modulation. The white contour indicates the contiguous cluster of affected voxels in aFST/IPa region. (**C,D**) Magnified versions of Figure 3A and 2B highlighting the most affected mid-STS aFST/IPa region during neglect in monkey # 1 (**C**) and monkey # 2 (**D**). (**E,F**) Coronal slices show the anatomical locations of the affected aFST/IPa region in monkey # 1 (**E**) and monkey # 2 (**F**). The white cross-hairs indicate the peaks of the aFST/IPa ROI from Figure 4A. See also Figure S2.

In the resulting F-statistic maps, a contiguous region in the mid-STS cortex stands out as showing the largest loss of attention-related modulation during neglect (Figure 3A,B). As clarified by a magnified view of the inflated mid-STS cortex, the affected voxels in this region spanned two anatomical areas – anterior FST and posterior IPa in both monkeys (Figure 3C,D); hence, we refer to this region as aFST/IPa. Coronal slices show that the effects were localized to the lower part of the floor of the STS in both monkeys (Figure 3E,F). The reductions were specific to the hemisphere on the same side as the SC inactivation (Monkey #1: F (1,24971) = −165.58; Monkey #2: F (1, 26645) = −111.59); in the other hemisphere, there were significant increases during neglect in both monkeys (Monkey #1: F (1,24971) = 29.83; Monkey #2: F (1, 26645) = 44.15).

To corroborate these results, we performed another, independent analysis by identifying regions of interest (ROIs), defined as 2mm-radius spheres centered on the local maxima of the attention-related modulation from the control sessions. Keeping these ROI definitions fixed, we then assessed how the modulation in these pre-defined ROIs changed during neglect. This analysis identified 9 ROIs, illustrated by the coronal slices and time-courses in Figure 4. We also computed d-prime values (which capture effects on both the mean and variance of the BOLD signal) to summarize the attention-related modulation during control and neglect (bar plots in Figure 4). We then compared the modulation (d’) during control and neglect session to identify ROIs with significant changes during neglect (bootstrap test; p<0.05; Bonferroni corrected), and the rows in Figure 4 are rank-ordered by the size of the decrease in attention-related modulation. Consistent with the voxel-wise ANOVA results (Figure 3), the largest reductions in attention-related modulation were in the ROI located in the mid-STS cortex and the effect was specific to the hemisphere ipsilateral to SC inactivation (Figure 4A,B). Smaller reductions were found in other cortical areas well-known to be involved in selective attention: there were significant reductions ipsilateral to SC inactivation in parts of prefrontal cortex (FEF) and parietal cortex (LIPd but not LIPv). There were also some changes in visual areas but the effects were not unilateral and were mixed across both monkeys.

**Figure 4.**
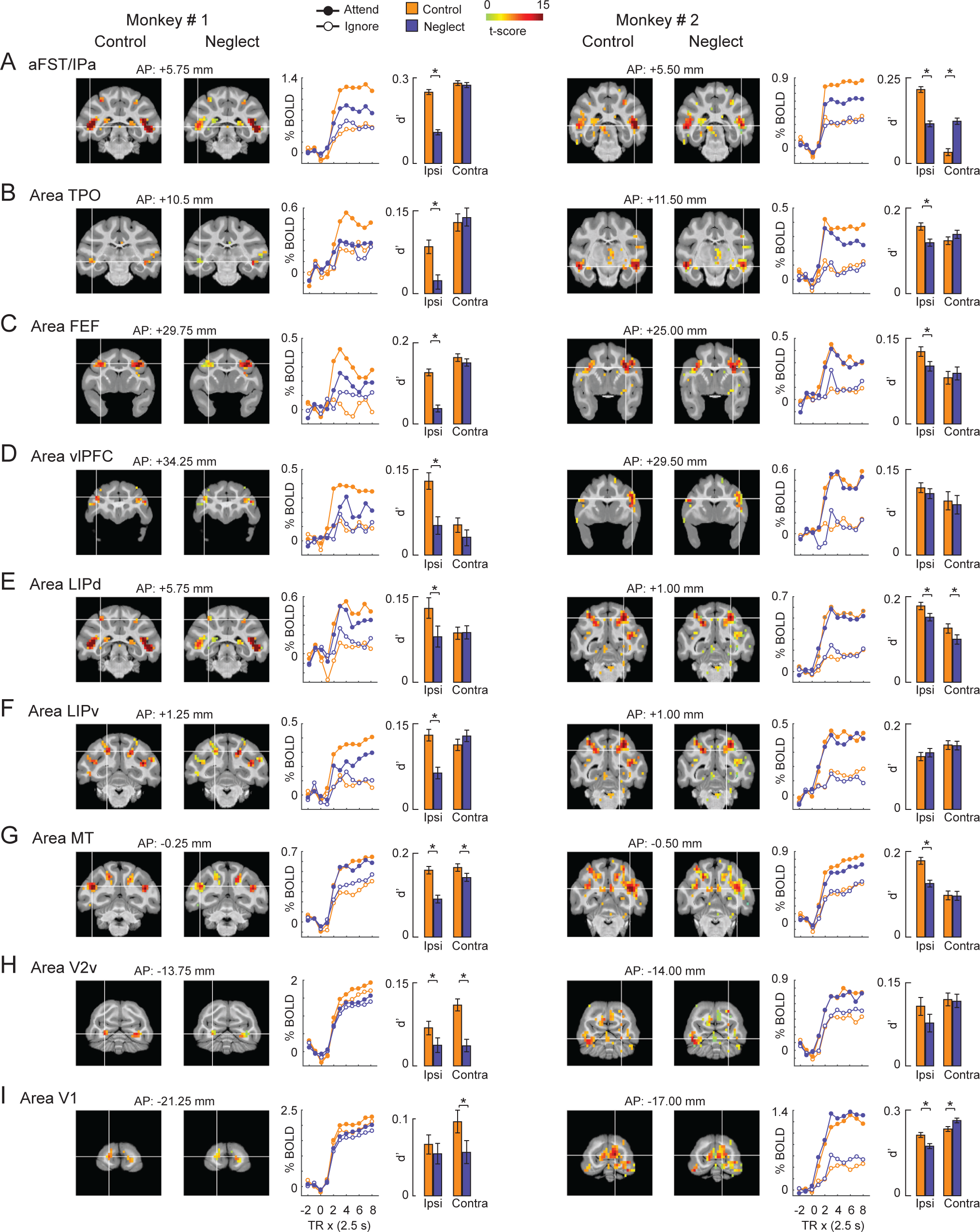
Effect of midbrain-induced neglect (SC inactivation) on the attention-related modulation in cortical ROIs. (**A-I**) The name of the ROI is shown at the top-left of each row. Each row shows coronal slices containing the peak of the ROI for both monkeys (Monkey # 1: Left column; Monkey # 2: Right column). The coronal slices overlaid with attention-related modulation during control and neglect (SC inactivation) sessions are shown next to each other for comparison. The location of the white cross-hair in each coronal slice indicates the peak of the ROI in the hemisphere ipsilateral to the SC inactivation site. Note that the peak of the ROI was identified based on the attention-related modulation during control. The location of the coronal slice w.r.t the inter-aural axis is shown on top of each coronal slice. Next to the coronal slices, mean BOLD time-courses for the ipsilateral ROI are plotted as % change in BOLD on y-axis against repetition time (TR) on the x-axis for the Attend (filled circles) and Ignore (open circles) tasks performed during control (orange) and neglect (blue) sessions. TR = 0 on x-axis indicates start of the block. The bar plot shows the attention-related modulation in the ipsilateral ROI measured as d’ during control (orange) and neglect (blue) sessions. For comparison, the same measure is shown for the corresponding contralateral ROI during control and neglect sessions. Error bars indicate bootstrapped 95% CI and * indicates statistical significance at 5% after correcting for multiple comparisons (p < 0.05; Bonferroni corrected). aFST refers to anterior part of anatomical area FST. See also Figures S3 and S4.

Direct comparison of these ROIs during neglect and control conditions illustrates the distinctive changes in attention-related BOLD modulation we found most prominently in the aFST/IPa region (Figure S3). The ROIs ipsilateral to SC inactivation generally all had lower d’ values during neglect compared to control (Figure S3A,C), but the largest reductions were found in aFST/IPa – the aFST/IPa is furthest from the line of unity slope (red arrows in Figure S3A,C). These reductions in attention-related BOLD modulation were spatially specific in aFST/IPa and some, but not all, of the other ROIs contralateral to SC inactivation (Figure S3B,D), as shown by the data points lying along or above the line of unity slope.

We performed several controls to validate these findings. Analysis of sessions with injections of saline versus muscimol into the SC confirm the findings that the mid-STS cortex, along with FEF, showed a unilateral reduction in modulation during midbrain-induced neglect (Figure S4). We verified that these findings did not depend on our choice of attention-related metric: we obtained the same result when we used the β coefficient of the term contrasting Attend and Ignore conditions – namely, the difference between mean BOLD in Attend and mean BOLD in Ignore (*μ*_*A*_-*μ*_*I*_). Taken together, these results demonstrate that an attention-related area in the mid-STS cortex (aFST/IPa) is causally implicated in visual spatial neglect.

### Mid-STS cortex selectively affected with static as well as dynamic stimuli

We next tested whether the loss of attention-related modulation in mid-STS cortex was specific to the use of visual motion stimuli in our attention tasks. We acquired additional fMRI sessions using the same task design, except that the motion stimuli were replaced with static white noise stimuli and motion-change detection was replaced with a second-order orientation detection (Figure 5A,B). We collected 7 no-injection (824 blocks each for both the Ignore and Attend tasks) and 9 SC inactivation sessions (848 blocks each for both the Ignore and Attend tasks), and confirmed that SC inactivation significantly affected performance for the contralateral second-order orientation changes compared to the ipsilateral second-order orientation changes in the Attend task during every session (Figure 5C; Chi-square proportion test; p<0.05).

**Figure 5.**
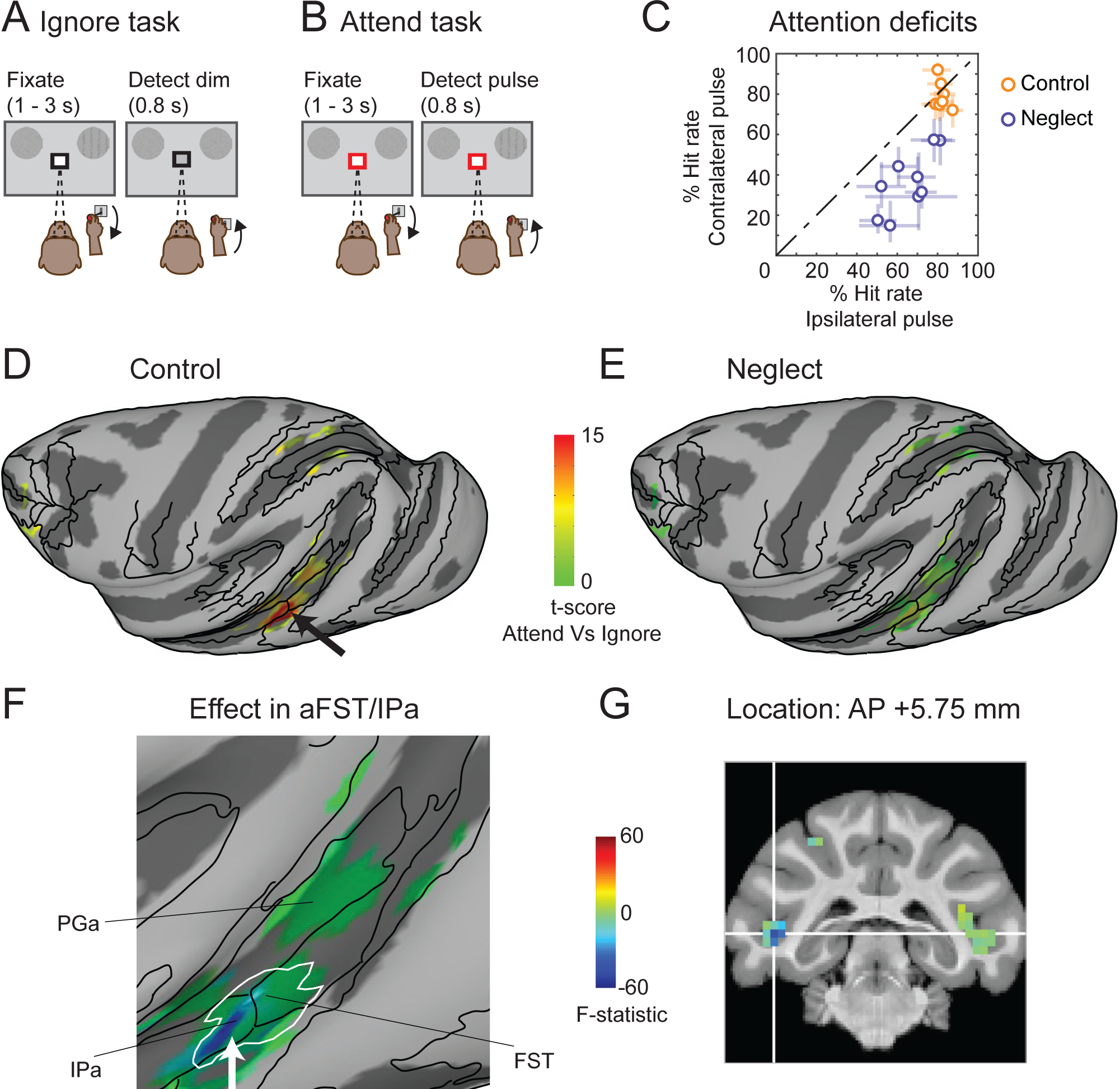
Effect in mid-STS cortex during neglect was not contingent on visual motion. (**A,B**) Same conventions as Figure 1B,C except that peripheral motion stimuli were replaced by white noise distractors and the motion-change event was replaced by a brief (0.5 s) second-order orientation pulse. (**C**) Performance in the Attend task is shown for control (green) and neglect (blue) sessions in monkey # 1. Hit rate (%) for orientation pulse in the affected visual field (y-axis) is plotted against hit rate for orientation pulse in the unaffected visual field (x-axis). Each data point represents a single session and error bars indicate 95% CI. (**D,E**) Inflated cortical maps of t-scores contrasting the Attend (Figure 5B) and Ignore (Figure 5A) tasks show attention-related modulation in the ipsilateral (left) hemisphere of monkey # 1 during control (**D**) and neglect (**E**) sessions. The black arrow points to the modulation in aFST/IPa region during control (**D**). The maps were thresholded during control sessions (**D**) and the modulation for the same voxels is shown during the neglect sessions (**E**). (**F**) Magnified version of the inflated cortical map of the interaction term F-statistic shows the changes in attention-related modulation during neglect in ipsilateral STS regions including aFST/IPa of monkey # 1 in the second-order orientation task. The effect during neglect is shown for the same voxels that showed significant modulation during control sessions (Figure 5D). The blue color indicates voxels whose modulation was reduced the most during neglect and the green colors indicate voxels with no reduction in modulation. The white contour indicates the aFST/IPa region affected during neglect in the motion task and is the same as shown in Figure 3C. (**G**) Coronal slice displays the anatomical location of affected aFST/IPa region in monkey #1 in the second-order orientation task. The white cross-hair indicates voxel with peak attention-related modulation in aFST/IPa region during control. See also Figure S5.

We first performed a voxel-wise analysis contrasting the BOLD measurements between the Attend and Ignore tasks, as before, and found significant attention-related modulation (t-scores, Bonferroni corrected; p<0.05) in the mid-STS cortex (black arrow in Figure 5D), as well as areas of posterior STS, IPS and the frontal cortex. As expected, the attention-related modulation in the second-order orientation task was sparser, because the orientation stimulus lacked the continuous dynamic changes of the motion stimulus. These results show that the aFST/IPa region in mid-STS cortex was modulated during covert visual attention even in the absence of visual motion.

We then determined if the attention-related modulation during the second-order orientation task was also changed during midbrain-induced neglect (Figure 5E), using the same voxel-wise ANOVA analysis described in the earlier section. The voxels most affected during neglect were once again identified in the mid-STS region at the border between anterior FST and posterior IPa on the side associated with neglect (F (1, 23083) = −63.64; white arrow in Figure 5F). There was no effect in the other hemisphere (F (1, 23083) = 0.000419), and region-of-interest analysis confirmed the largest reduction in aFST/IPa (Figure S5A). The affected voxels in the mid-STS (blue region in Figure 5F) completely overlapped with the aFST/IPa region that was affected in the motion task (white contour in Figure 5F), in the same lower part of the fundus in the STS at the identical anterior-posterior location (compare Figure 5G and Figure 3E). These results demonstrate that the same region of mid-STS cortex is causally linked to neglect even when the task requires attention to second-order orientation rather than motion.

### Mid-STS cortex selectively affected during neglect caused by FEF inactivation

Because neglect is associated with both cortical and subcortical attention-related circuits, we next tested whether the same mid-STS region was affected by inactivation of frontal eye fields (FEF) in prefrontal cortex. We identified FEF in the anterior bank of the arcuate sulcus using microstimulation with currents of 40 μA to evoke saccades (Figure 6A). For reversible inactivation, we injected 1-2 μl of muscimol and mapped the affected visual field using a visually guided saccade task (Figure 6B), confirming that the affected portion of the visual field overlapped the contralateral stimulus in the attention task. During the attention task, we found a significant effect on the median response times for contralateral motion-changes compared to ipsilateral motion-changes (median RT_contra_ = 545 ms; RT_ipsi_ = 521 ms; Wilcoxon rank sum test; p<0.001), although hit rates were not significantly changed (Chi-square proportion test; p>0.05). Overall, we collected 7 no-injection (768 blocks each for both the Ignore and Attend tasks) and 7 inactivation sessions (736 blocks each for both the Ignore and Attend tasks).

**Figure 6.**
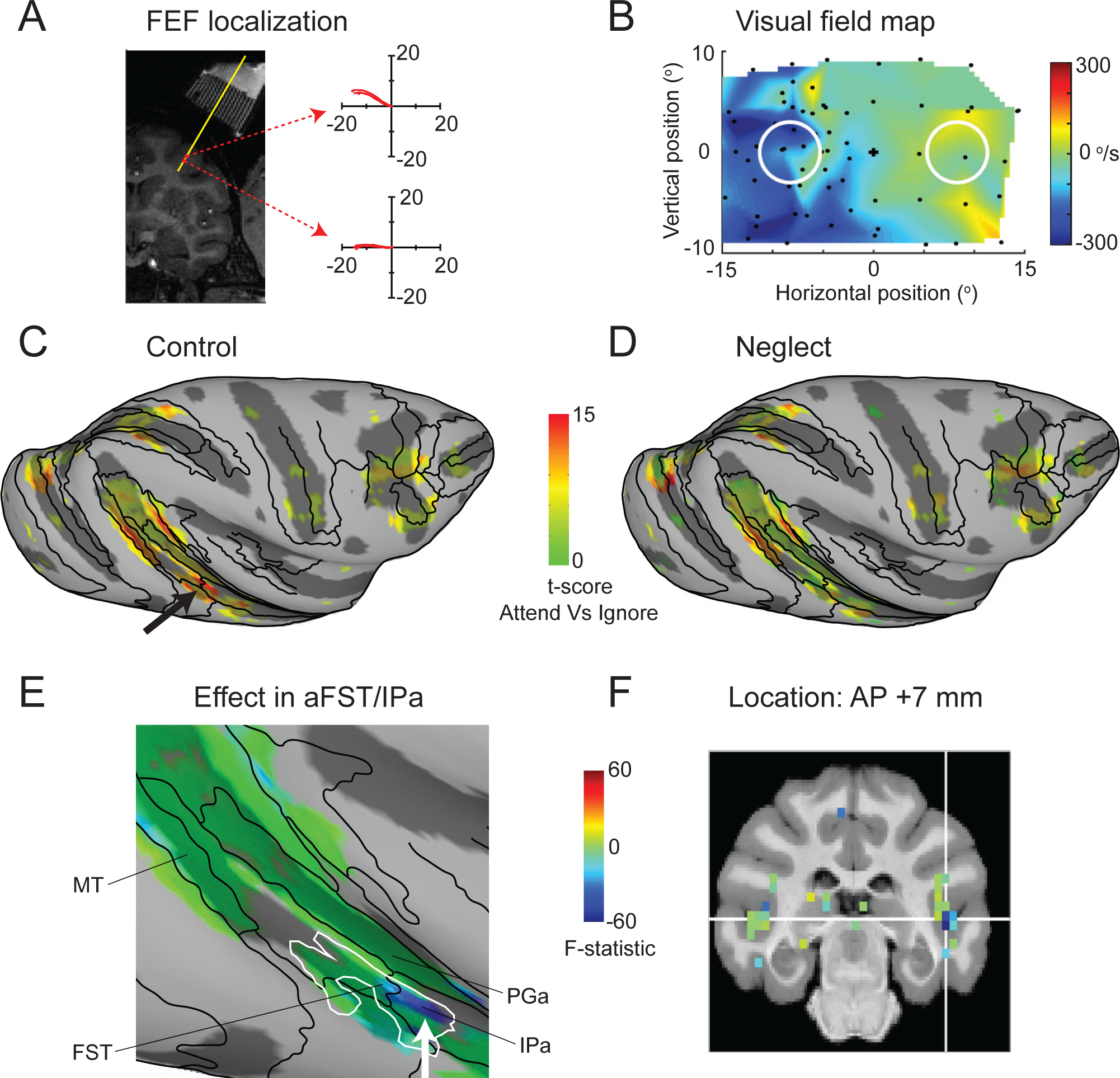
The same mid-STS region was affected during FEF-induced neglect. (**A**) Coronal section shows the localization of FEF in the anterior bank of arcuate sulcus and illustrates two sample sites (red circles) at which electrical stimulation (70 ms train duration, 350 Hz, biphasic pulses with duration of 0.3 ms, 40 μA) evoked saccadic eye movements with directions and amplitudes (indicated by the trajectories in each of the two corresponding panels) that depended on the location within the FEF. White stripes in upper right of the MRI were produced by the contrast agent placed in the FEF recording grid and the superimposed yellow line indicate the estimated electrode and injectrode paths. (**B**) The visual field map shows the affected visual field (blue region) during FEF inactivation. Same conventions as Figure 1E (top panel). (**C,D**) Inflated cortical maps of t-scores contrasting the Attend (Figure 1C) and Ignore (Figure 1B) tasks show attention-related modulation in the ipsilateral (right) hemisphere of monkey #2 during control (**C**) and neglect (**D**) sessions. The black arrow points to the modulation in aFST/IPa region during control (**C**). The maps were thresholded during control sessions (**C**) and the modulation for the same voxels is shown during the neglect sessions (**D**). **(E)** Magnified version of the inflated cortical map of the interaction term F-statistic shows the changes in attention-related modulation during FEF-induced neglect in ipsilateral STS regions including aFST/IPa of monkey #2 in the motion task. The effect during neglect is shown for the same voxels that showed significant modulation during control sessions (Figure 6C). The blue color indicates voxels whose modulation was reduced the most during neglect and the green colors indicate voxels with no reduction in modulation. The white contour indicates the aFST/IPa region affected during midbrain-induced neglect (SC inactivation) in motion task and is the same as shown in Figure 3D. **(F)** Coronal slice shows the anatomical location of the affected aFST/IPa region during FEF inactivation in monkey #1. The white cross-hair indicates voxel with peak attention-related modulation in aFST/IPa region during control sessions. See also Figure S5.

As expected, during control sessions there was significant attention-related modulation in several cortical areas including the mid-STS region (Figure 6C). During FEF inactivation, the most strongly affected voxels were again found in the ipsilateral mid-STS region at the border between anterior FST and IPa (F (1, 21322) = −93; white arrow in Figure 6E, see also Figure S5B), with no significant effect in the contralateral mid-STS region (F (1, 21322) = 2.684). Once again, the region identified during FEF inactivation (blue region) fully overlapped with the region identified during midbrain-induced neglect (white contour). Region-of-interest analysis showed that the affected region was localized to the fundus of the STS at nearly the same anterior-posterior location as that identified during midbrain-induced neglect (Figure 6F). Although the size of the changes during FEF inactivation were smaller than those during superior colliculus inactivation, consistent with the weaker effects of inactivation on attention task performance [41], these results demonstrate that the same mid-STS region identified during midbrain-induced neglect is also causally linked to prefrontal attention areas.

### Reversible inactivation of the mid-STS region itself caused neglect

Finally, given the conspicuous drop in attention-related modulation in the mid-STS region during both midbrain-and prefrontal-induced neglect, we tested the behavioral consequence of directly inactivating this region itself. Following procedures similar to those used during inactivation of the SC and FEF, we injected 4-6 μl of muscimol into the center of the aFST/IPa region identified in monkey #1, guided by the structural MRI localization of the ROI.

Unlike SC or FEF, reversible inactivation of this mid-STS region had no detectable effects on the metrics of saccadic eye movements (Figure 7A) demonstrating that the unilateral effects in this region during midbrain-and prefrontal-induced neglect were not related to suppression or facilitation of eye movements. However, inactivation dramatically impaired detection performance in the Attend task for contralateral stimulus changes (Figure 7B), causing neglect-like deficits at least as large as those observed during inactivation of the SC (Figure 1F). During all 6 inactivation sessions, there was a significant reduction in hit rate for motion-direction changes for the stimulus in the contralateral visual field compared to the stimulus in the ipsilateral visual field (Chi-square proportion test; p<0.05); there was no significant difference in hit rates for the no-injection and saline-injection control sessions (orange and black symbols).

**Figure 7.**
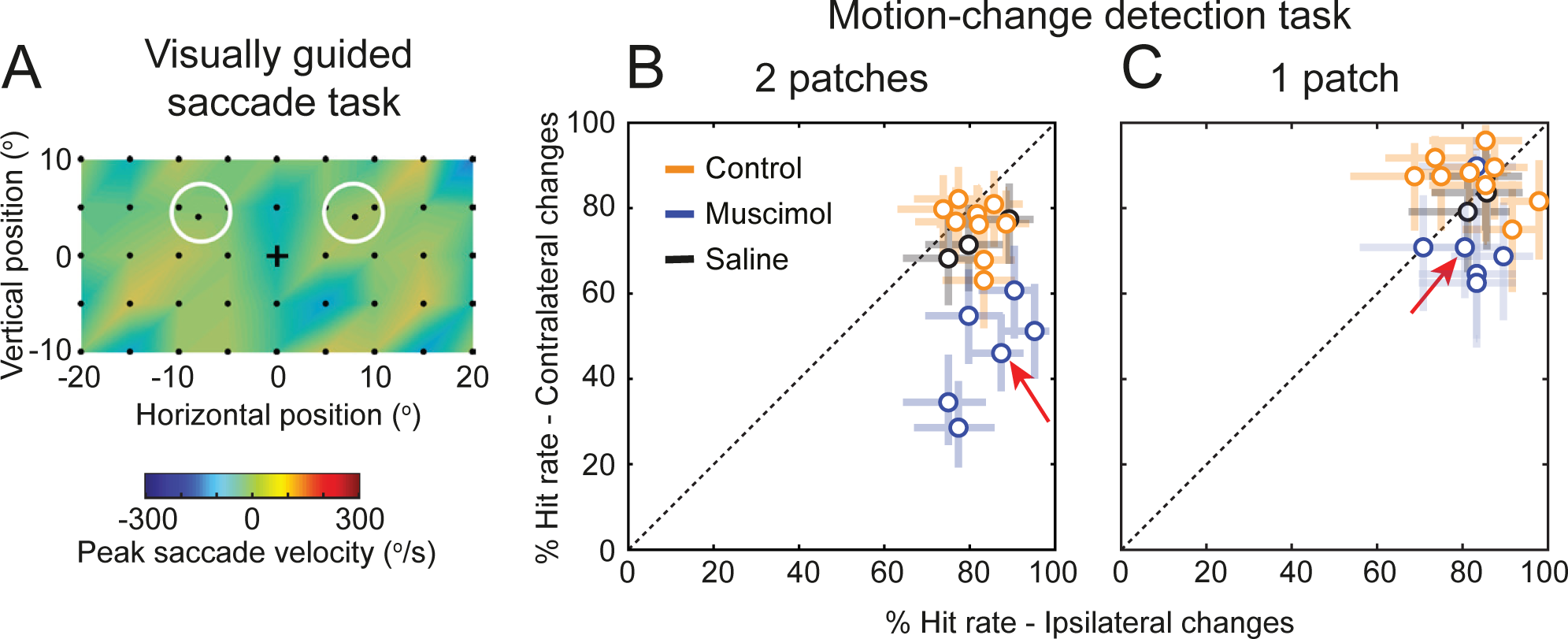
Reversible inactivation of the mid-STS region causes neglect-like deficits but does not affect delayed visually guided saccades. The aFST/IPa region in monkey #1 was reversibly inactivated using 4-6 ul of muscimol. Following the inactivation, we tested the behavior of the monkey in the same delayed visually guided saccade task and Attend task that we used for SC inactivation experiments (Figure 1E,F). (**A**) The visual field map shows the effect of aFST/IPa inactivation on the peak velocity of the saccades to targets in the visually guided saccade task, from 1 sample inactivation session. Decreases in peak velocity would appear as blue regions. Unlike what was observed after SC inactivation (Figure 1E), inactivation of aFST/IPa caused no evident changes in saccade velocities in any of the 6 sessions. Black dots indicate the retinotopic position of the saccade targets. (**B**) Performance in Attend task (with two motion patches) is shown for no-injection controls (orange), muscimol injection (blue), and saline injection (black) sessions. % Hit rate for motion changes in the affected visual field (y-axis) is plotted against % hit rate for motion changes in the unaffected visual field (x-axis). Each data point represents a single session and error bars indicate 95% CI. Significant differences in hit rate were found in all 6 inactivation sessions. Red arrow indicates the data point from the same session illustrated in panel (**A**). (**C**) Performance in the Attend task during the control condition in which only a single patch was presented. Same conventions as in panel (**B**).

Moreover, the disruption in detection performance was markedly reduced when only a single motion patch was presented on each trial (Figure 7C), demonstrating that the deficits had extinction-like properties [42] consistent with spatial neglect. These results directly demonstrate that this circumscribed region of mid-STS cortex plays a causal role in the covert orienting of attention – the central functions that are disrupted during spatial neglect – without affecting overt aspects of orienting such as the generation or suppression of saccadic eye movements.

## Discussion

Using fMRI and reversible inactivation in monkeys performing attention tasks, we identified a localized region in the floor of the mid-STS cortex (aFST/IPa) as the most strongly affected patch of cortex during visual spatial neglect. Several aspects of our results point to the uniqueness of this patch of cortex. The loss of attention-related modulation did not depend on the specific visual feature attended, indicating a general role in visual selection rather than processing of particular features. The same region was affected by neglect induced by inactivation of the SC in the midbrain and the FEF in prefrontal cortex, showing that it is a shared node in both cortical and subcortical attention circuits. Moreover, focal inactivation of this region itself caused neglect-like deficits in covert attention, but without affecting saccadic eye movements, unlike other brain regions in monkey implicated in control of attention (e.g., SC, FEF). Together, our results demonstrate for the first time a causal link between spatial neglect and an attention-related region in the mid-STS cortex of monkeys.

It may seem surprising that our results single out a region in mid-STS cortex, because most studies of attentional control in nonhuman primates have focused on the roles of frontal and parietal cortex [43,44]. However, recent fMRI studies in monkeys have identified attention-related regions in the superior temporal sulcus, as well as in frontal and parietal areas [33-35,39]. A similar mid-STS region, albeit in the upper bank of STS, was also identified in monkeys performing memory-guided saccades, before and during reversible inactivation of area LIP, suggesting this region was involved in compensating for the orienting deficits caused by parietal inactivation [45]. More specifically, a region in the posterior inferotemporal cortex (referred to as PITd) is strongly modulated during attention to motion stimuli, at similar anterior-posterior locations in the STS as the region we identified as aFST/IPa; the attention-related activation in this mid-STS region is not explained by eye movements, because it occurs during covertly performed tasks, and does not depend on the specific visual feature attended [34,35]. By showing that this mid-STS region is causally linked to other major attention-related structures – and is itself necessary for the normal control of covert attention – our results provide direct evidence that the cortical circuits for attention in monkeys include a crucial node in the mid-STS cortex.

The localization of this region in the mid-STS cortex in monkeys is broadly consistent with results from human studies. The anatomy of hemi-spatial neglect includes regions of the superior temporal cortex, especially the temporoparietal junction, that lies at the posterior end of the superior temporal sulcus in humans [6,25]. This temporal cortical region is also a key node in the ventral attention network, which is implicated in interrupting the current focus of attention in favor of a novel object of interest [46]. The mid-STS region in monkeys provides a new experimental avenue for investigating how the neuronal processing in this node contributes to the control of attention, and specifically, how disruption of these processing steps causes the symptoms of spatial neglect. It is probably not a coincidence that this mid-STS region is also a major site of convergence between the dorsal “where” and ventral “what” streams of visual processing in macaques [47,48].

However, homology is notoriously difficult to establish, and there are major differences in the organization of these areas in humans and monkeys. The mid-STS region in monkeys has a similar functional connectivity to the temporoparietal junction area in humans [49], but the mid-STS region is notably not activated during the same rapid serial visual presentation task that reliably identifies the TPJ area in humans [36]. More fundamentally, spatial neglect in humans is predominantly associated with right hemisphere lesions [3,6], whereas the neglect-like deficits generated in monkeys, here and previously, are equally well achieved with inactivation on either side of the brain [28-32]. The strong lateralization of cortical functions in humans is a basic difference from monkeys that will require teasing apart functions that are shared versus specialized across primate species.

## Acknowledgments

We thank Tom Ruffner and Nick Nichols for technical support and Fabrice Arcizet, James Herman, Leor Katz and Lupeng Wang for helpful input. We also thank Brian Russ for helpful suggestions in setting up fMRI preprocessing pipeline and constructing inflated maps. This work was supported by the National Eye Institute Intramural Research Program at the National Institutes of Health. Functional and anatomical MRI scanning was carried out in the Neurophysiology Imaging Facility Core (NIMH, NINDS, NEI) supported under IRP grant ZIC MH002899. This work utilized the computational resources of the NIH HPC Biowulf cluster.

## Author Contributions

All authors designed the experiments. A.R.B. and A.B. conducted experiments and collected the data. A.R.B. analyzed the data. All authors interpreted the results. A.R.B. and R.J.K wrote the manuscript.

## STAR Methods

### CONTACT FOR REAGENT AND RESOURCE SHARING

Further information and requests for resources should be directed to and will be fulfilled by the Lead Contact: Richard J. Krauzlis

### EXPERIMENTAL MODEL AND SUBJECT DETAILS

#### Animals

Two adult male rhesus monkeys (*Macaca mulatta*) weighing 7-9 kg participated in this study. All experimental protocols were approved by the National Eye Institute Animal Care and Use Committee and all procedures were performed in accordance with the United States Public Health Service policy on the humane care and use of laboratory animals. Under isoflurane and aseptic conditions, we surgically implanted plastic head-posts and electrophysiology chambers to access the SC and FEF. The SC chamber was angled 38° to the posterior of vertical and directed at the midline, 15 mm above and 1 mm posterior to the interaural line. The FEF chamber was angled 30° lateral of vertical and aimed at a point 18 mm lateral from the midline, 25 mm anterior to the interaural line.

### METHOD DETAILS

#### Experimental apparatus

Monkeys were seated and head-fixed in a custom-built MR-safe chair with a joystick attached inside the chair. Before each imaging session, monkeys performed a visual guided saccade task inside a darkened booth. The saccade task was controlled using a modified version of PLDAPS [1]. Animals viewed visual stimuli from a distance of 48 cm that were displayed at 1920 x 1200 resolution (∼ 60° x 38°) and 100Hz frame-rate on a VIEWPixx display (VPixx Technologies, Saint-Bruno, QC Canada), controlled by a mid-2010 Mac Pro (Apple Inc., Cupertino, CA) running MATLAB (The Mathworks, Natick, MA) with the Psychophysics Toolbox extensions [2-4]. Eye position was recorded using an EyeLink 1000 infrared eye-tracking system (SR Research Ltd., Ottowa, Ontario, Canada).

During each imaging session, monkeys performed attention tasks and visual guided saccade task inside the vertical scanner. Stimuli were back projected on to a screen placed inside the bore of the vertical scanner using an Epson projector controlled by a Windows 2007 machine running MATLAB R2012b (The Mathworks) with the psychophysics toolbox extensions [2]. The timing of the stimuli and events were controlled by a QNX system running QPCS. Monkey viewed the screen through a mirror placed in front at a 45° angle. The total viewing distance of the screen was 53 cm. Eye movements were acquired and monitored in the scanner using an iView system (Version 2.4, SensoMotoric Instruments). Eye signal was calibrated at the beginning of each session. Joystick presses and releases were detected by disruption of an optic-fiber transmission using a custom-built device. Joystick was calibrated once at the beginning of the experiments.

#### fMRI data collection

Anatomical and functional images were collected in a vertical magnet (4.7T, 60 cm vertical bore; Bruker Biospec) equipped with a Bruker S380 gradient coil. EPI volumes were acquired using a custom built transmit and 4-channel receive RF coil system (Rapid MR International). In each run, we collected 192 whole-brain EPI volumes at an isotropic voxel resolution of 1.5 mm and at a TR of 2.5 s.

#### Attention tasks: Motion-change detection

In the scanner, both monkeys performed three behavioral tasks: Baseline, Ignore and Attend. In all tasks, monkeys initiated the trial by holding the joystick down and fixating the central fixation spot with a colored central cue on a grey background. Monkeys fixated for the entire duration of the trial with in a 2° fixation window. The color of the central cue indicated the trial condition. In Baseline and Ignore trials, the color of the central cue was black (Figure 1A, 1B) and the relevant stimulus was the fixation stimulus. In Attend trials, the color of the central cue was red (Figure 1C) and the relevant stimulus was the peripheral motion stimulus. The sequence of events in three different trial conditions are as follows.

In Baseline trials, following 0.5 s of fixation, the luminance of the fixation spot decreased during a variable delay of 1 – 3 s on the 65% of the trials. Monkeys reported the luminance change by releasing the joystick within 0.3 −0.8 s to get a juice reward (Figure 1A). A total of 8401 trials (4433 in Monkey #1 and 3968 in Monkey #2) were collected during control and 7484 trials (3667 in Monkey #1 and 3817 in Monkey #2) were collected during SC inactivation in both monkeys. We also collected 4446 trials during saline injection in monkey #1.

In Ignore trials, following 0.5 s of fixation, two random dot motion stimuli were presented on either side of fixation at 8° eccentricity (radius) and 10° above horizontal meridian (azimuth). During the variable delay of 1 – 3 s, the luminance of the fixation spot decreased on 65% of the trials. Independent of the fixation luminance change, one of the peripheral motion stimuli changed direction during the variable delay of 1 – 3 s on the 65% of the trials. Monkeys ignored the motion direction change and reported the luminance change by releasing the joystick within 0.3 - 0.8 s to get a juice reward (Figure 1C). If the monkeys released the joystick for a motion direction change, the trial was aborted. A total of 9139 trials (4912 in Monkey #1 and 4227 in Monkey #2) were collected during control and 8240 trials (4083 in Monkey #1 and 4157 in Monkey #2) were collected during SC inactivation in both monkeys. We also collected 4913 trials during saline injection in monkey #1.

In Attend trials, following 0.5 s of fixation, two random dot motion stimuli were presented at the same stimulus location as in Ignore trials. One of the peripheral motion stimuli changed direction during the variable delay of 1 – 3 s on the 65% of the trials. Monkeys reported the motion-direction change by releasing the joystick within 0.3 - 0.8 s to get a juice reward (Figure 1E). There was no fixation luminance change in Attend trials. A total of 8672 trials (4765 in Monkey #1 and 3907 in Monkey #2) were collected during control and 7742 trials (4026 in Monkey #1 and 3716 in Monkey #2) were collected during SC inactivation in both monkeys. We also collected 4794 trials during saline injection in monkey #1.

In all tasks, 35% of the trials were catch trials and monkeys hold the joystick down to get a juice reward.

#### Random dot motion stimuli

The random dot motion stimuli were circular patches of moving dots, with the direction of motion of each dot drawn from a normal distribution with a mean value (defined as the patch motion direction) at 30° above horizontal and a 16° standard deviation. The lifetime (10 frames, 100 ms), density (25 dots/°^2^/s), and speed of the dots (15 °/s) were held constant. The radius of the aperture was set to 3°. Luminance of each moving dot in the motion patches was 50 cd/m^2^. The change in direction of motion (Δ) was 1 ± 0.25 standard deviations for both monkeys across sessions.

#### Fixation spot stimulus

The size of the fixation spot was 0.23° and the size of the central cue was 0.35°. The background luminance of the screen was 14 cd/m^2^ and the luminance of the fixation spot was 50 cd/m^2^. The luminance change in fixation spot during Baseline and Ignore trials was 1-2 cd/m^2^ across sessions for both monkeys.

#### Attention tasks: Orientation-pulse detection

In addition to the motion-change detection task, monkey # 1 also performed a version of the attention tasks with orientation pulse stimuli instead of the random dot motion stimuli. The sequence of events in all three conditions (Baseline, Ignore, Attend) was kept the same as the motion-change detection version of the task. The onset of the motion stimuli was replaced with the onset of white noise stimuli, and the motion-direction change was replaced with a 0.5 s second-order orientation pulse (Figure 6A, 6B). In the Attend condition, the monkey reported the orientation pulse by releasing the joystick within 0.3 - 0.8 s to get a juice reward (Figure 6A), whereas in the Ignore condition, monkey ignored the orientation pulse and reported the luminance change in the fixation spot by releasing the joystick within 0.3 - 0.8 s to get a juice reward (Figure 6B). In the Attend condition, a total of 3700 trials were collected during control and 2680 trials were collected during SC inactivation. In the Ignore condition, a total of 3752 trials were collected during control and 2698 trials were collected during SC inactivation.

The second-order orientation stimulus was generated by briefly (0.5 s) modulating the contrast of a white noise stimulus with a 2-dimensional sinusoid. The noise stimulus was 6^0^ in diameter and consisted of checks each the size of a pixel with luminance values ranging from 8 – 84 cd/m^2^, and the 2-dimensional sinusoid had a spatial frequency of 0.7 cycles/deg, and its orientation was 90°. We refer to this as a second-order orientation stimulus, because the oriented grating briefly visible in the stimulus was due to the local differences in contrast, not luminance differences. The mean luminance (38 cd/m^2^) of the stimulus was constant throughout its presentation and was the same across every band in the oriented grating.

#### Block Design

Baseline, Ignore and Attend tasks were presented in a block design as shown in Figure 1D. Each run started with a Baseline block which lasted for 10 s and was presented in every alternate block thereafter. Ignore and Attend blocks lasted for 20 s and were presented randomly between Baseline blocks. The number of Ignore and Attend blocks were balanced in a given run. Each run lasted 480s.

#### Electrophysiology

Locations for muscimol injections in the SC and FEF were identified by single-unit recording and electrical microstimulation. Single-unit activity was recorded using tungsten in glass-coated electrodes with impedances of 1-2 MOhm (Alpha Omega Co., Inc., Alpharetta, GA). Electrode position was controlled with a Narishige microdrive. The electrical signal was amplified and recorded online using the OmniPlex system (Plexon Inc., Dallas, Texas). Response fields and saccade-related movement fields were mapped by having the monkey perform the visually guided saccade task; we confirmed saccade-related activity consistent with the known activity patterns in the FEF or SC. Electrical microstimulation was also applied (70 ms train duration, 350 Hz, biphasic pulses with duration of 0.25 ms) to evoke saccades with currents (40 μA in FEF; 20 μA in SC) confirming that we were in the FEF or intermediate layers in SC [5,6]. All candidate sites were first identified by neuronal recordings and electrical stimulation prior to the muscimol inactivation experiment; in some cases, the FEF or SC site was verified immediately prior to muscimol injection using the electrode placed within the injection cannula [7].

#### Muscimol injections

Reversible inactivation of the intermediate layers of the SC (n = 27; monkey #1(LH): 19, monkey #2 (RH): 8), area FEF (n = 7; monkey #2 (RH)), and the aFST/IPa region (n = 6; monkey #1 (LH)) were done by injecting muscimol (5 mg/ml) in either the left (LH) or right (RH) hemisphere of each monkey. The amount of muscimol injected ranged from 0.3-0.5 μl in SC, 1-2 μl in FEF, and 4-6 μl in aFST/IPa. Injections were done using a custom-made apparatus modified from [8], with an injection pump at a rate of 0.5 μl in 10 mins. For saline controls, the same volumes of saline were injected at the same locations in the intermediate layers of the SC or in the aFST/IPa region.

#### Visually guided saccade task and mapping of affected visual field

In the saccade task, each trial started with the monkeys fixating a small square stimulus (0.23° wide, 50 cd/m^2^) placed at the center of the visual display. After 500 ms of fixation, a second target (0.23° wide, 50 cd/m^2^) was presented at some other location in the visual display. Monkeys were required to maintain fixation until the central fixation stimulus was turned off, 1-2 seconds after the onset of the peripheral target. At that point, the monkeys should make a saccade to the second, peripheral target and fixate it for at least 500 ms in order to receive a reward. The eye position data were used to detect saccades and quantify saccade metrics offline using methods described previously [9].

Before imaging, monkeys performed the saccade task outside the scanner and the location of the second target was systematically varied every trial in order to quantify the metrics of saccades to targets across the visual field. We quantified the impairments in saccade metrics during SC or FEF inactivation by measuring the difference in the peak velocity for saccades made to targets at different locations in the visual field before and during inactivation. The target locations where the peak velocities of the saccades were impaired identified the affected visual field (Figure 1E, 6B). Data were collected beginning 30 minutes after the end of the muscimol infusion.

Inside the scanner, monkeys performed the saccade task every 8 minutes and the location of the second target was alternated each trial between contralateral and ipsilateral stimulus locations (radius: 8°; azimuth: 10°). The time course of muscimol effect was measured as a difference in the saccade latency to targets in the contralateral affected visual field compared to targets in the ipsilateral unaffected visual field (Figure 1E). We measured the impairment on the saccade latency instead of peak velocity in the scanner because, the low sampling frequency of the eye tracker in the scanner limited the accurate measurement of peak velocities.

#### Stimulus mapping experiment

To identify voxels responding to foveal and peripheral stimuli locations, a flickering checker board stimulus (4 Hz) that had concentric rings of 2° width spanning up to 12° eccentricity was used. In foveal visual stimulation blocks, the checker board stimulus was masked everywhere except for the central 2° radius. In peripheral visual stimulation blocks, the checker board stimulus was masked everywhere except for the two eccentric stimulus locations used in the attention tasks. The foveal and peripheral stimulation blocks (20 s duration) were interleaved with fixation blocks (10 s duration). A total of 258 runs (150 in Monkey #1; 108 in Monkey # 2) were collected in both monkeys across 13 sessions. Functional maps showing peripheral and foveal voxels were created using the same methods as described for creating functional maps for the attention tasks.

### QUANTIFICATION AND STATISTICAL ANALYSIS

#### fMRI Analysis

Preprocessing of the fMRI data was done using AFNI/SUMA software package [10]. Raw images were converted to AFNI data format. EPI volumes in each run were slice-time corrected using *3dTshift*, followed by correction for static magnetic field inhomogeneities using the *PLACE* algorithm [11]. All EPI volumes were motion-corrected using *3dvolreg* and were registered to the session anatomical using *3dAllineate*. To combine EPI data across multiple sessions for a given animal, all sessions for a given animal were registered to a template session. A high resolution anatomical of each monkey was registered to the corresponding template session anatomical to overlay functional results in monkey’s native space. To overlay D99 atlas boundaries on the functional results, D99 anatomical in AFNI was registered to each monkey’s native space [12]. Surface maps were generated for D99 anatomical in each monkey’s native space using CARET from a white matter segmentation mask [13]. Surface maps of D99 in each monkey’s native space were viewed in SUMA overlaid with anatomical boundaries and functional results.

#### Functional maps of attention-related modulation

To identify voxels that were modulated by attention we performed a GLM analysis using *3dDeconvolve* in AFNI. Attend and Ignore conditions were included as regressors of interest and Baseline condition, motion correction parameters were included as regressors of no interest (baseline model). To control for the effects caused by any differences during Attend and Ignore blocks in blinks (Wilcoxon rank sum test, p = 0.007 (Monkey # 1), p = 0.99 (Monkey # 2)), saccadic eye movements within the fixation window (Wilcoxon rank sum test, p < 0.0001 (Monkey # 1), p < 0.0001 (Monkey # 2)), rewards (Wilcoxon rank sum test, p < 0.0001 (Monkey # 1), p < 0.0001 (Monkey # 2)) and joystick movements (Wilcoxon rank sum test, p = 0.79 (Monkey # 1), p = 0.53 (Monkey # 2)), we included these factors as part of the regression model. Reward times, joystick event times (press and release), blink times and saccade times (left and right) were convolved with the hemodynamic impulse response function and were included as part of the baseline model. In the blocks of all three conditions for both monkeys, the median duration of fixation was above 85% of the block duration. Saccades within the fixation window (magnitude less than 1°) were detected using previously described methods [9].

T-scores contrasting Attend and Ignore conditions were projected on to the inflated maps to show voxels modulated by attention. All functional maps were thresholded (t-score > 5.02) to correct for multiple comparisons (p<0.05, Bonferroni correction). Voxels that mapped onto foveal locations, based on a separate retinotopic mapping experiment (described below), were excluded from the inflated maps in order to focus on BOLD changes related to whether or not the peripheral motion stimuli were attended, rather than the very small stimulus differences at the fovea in the two conditions (black versus red cue; Figure 1B,C).

#### Functional regions of interest and attention-related modulation

Functional regions of interest (ROIs) were identified based on the local maxima of the attention-related modulation map from the control sessions for each hemisphere using *3dExtrema* in AFNI. The BOLD time-course for each Attend or Ignore block was computed as a % change in BOLD relative to the BOLD in Baseline block preceding it. For each ROI, an average time-course was constructed by pooling time-courses of all voxels within a 2mm radius around the local maxima across all blocks from all sessions. D-prime was computed for each ROI as a measure of attention-related modulation from the Attend and Ignore distributions of BOLD values taken from the TRs 2 – 8 of each block. Attend and Ignore distributions were constructed by pooling BOLD values across voxels in a given ROI and across blocks from different sessions. The mean and variance of the Attend 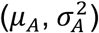 and Ignore 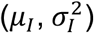 distributions were used to compute d-prime (*d*′) following equation 1. The source of the variance in the Attend and Ignore distributions came from pooling BOLD values across blocks, sessions and voxels in a given ROI.

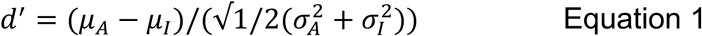

#### Voxel-wise ANOVA

For each voxel, we performed ANOVA with two factors and two levels each – covert attention (Attend and Ignore conditions) and neglect (control and inactivation). The 2×2 factorial design for each voxel was based on the BOLD values taken from the TRs 2 – 8 of Attend and Ignore time courses extracted from the control and neglect sessions. F-statistic of the interaction term was projected onto the inflated maps to show the effect of inactivation on attention-related modulation. For every voxel, the sign of the difference in attention-related modulation between neglect and control (*d*′_*Neglect*_-*d*′_*Control*_) was added to the F-statistic of the interaction term to differentiate the decrease in modulation from increase in modulation.

## Supplemental Information Titles and Legends

**Figure S1. Cortical maps of attention-related modulation in the contralateral hemisphere during control and neglect sessions (related to Figure 2)**

(**A,B**) Inflated cortical maps of t-scores showing attention-related modulation in contralateral (right) hemisphere of monkey # 1 during control (**A**) and neglect sessions (**B**). The maps were thresholded during control sessions (**A**) and the modulation for the same voxels is shown during the neglect sessions (**B**).

(**C,D**) Same conventions as (**A,B**) but for monkey # 2.

**Figure S2. Effect of midbrain-induced neglect (SC inactivation) on all voxels and voxels that map onto foveal locations (related to Figure 3)**

Inflated cortical maps of interaction term F-statistic show voxels whose modulation was affected during neglect induced by SC inactivation in the ipsilateral hemispheres of both monkeys. The blue color indicates voxels whose modulation was reduced the most during neglect and the green colors indicate voxels with no reduction in modulation. (**A,C**) The effect during neglect is shown for all voxels in the ipsilateral hemisphere of monkey # 1 (**A**) and monkey # 2 (**C**). (**B,D**) The effect during neglect is shown for voxels that map onto foveal locations in the ipsilateral hemisphere of monkey # 1 (**B**) and monkey # 2 (**D**). Cortical regions corresponding to the fovea were unaffected by midbrain-induced neglect following SC inactivation; the main effect of SC inactivation was diminished attention-related enhancement in *non-foveal* regions.

**Figure S3. Comparison of the effects of midbrain-induced neglect on the attention-related modulations in cortical ROIs (related to Figure 4)**

For both monkeys, SC inactivation had the strongest effects on the ipsilateral aFST/IPa region (red arrows). Attention-related modulation during neglect sessions is plotted against the attention-related modulation during control sessions for each ROI in the ipsilateral (**A,C**) and contralateral (**B,D**) hemispheres of both monkeys. In ROIs that lie significantly below the line of identity, attention-related modulation during neglect was significantly reduced compared to the modulation during control. The error bars indicate boot-strapped 95% confidence intervals. Among all ROIs in the ipsilateral hemispheres of monkey #1 (**A**) and monkey #2 (**C**), the aFST/IPa ROI (red symbol and arrows) showed the strongest reduction in modulation and hence lies the furthest from the line of unity slope. In contrast, the aFST/IPa regions in the contralateral hemisphere of monkey #1 (**B**) and monkey #2 (**D**) did not show significant reductions.

**Figure S4. Comparison of attention-related modulation in cortical ROIs between neglect (SC inactivation) and saline sessions (related to Figure 4)**

(**A,B**) Inflated cortical maps of t-scores showing attention-related modulation in ipsilateral (**A**) and contralateral (**B**) hemispheres of monkey # 1 during saline injection sessions. The maps were thresholded during control sessions (Figure 2, Figure S1) and the modulation for the same voxels is shown during the saline sessions (**A,B**).

(**C,D**) In the ipsilateral and contralateral ROIs defined based on the attention-related modulation map during control (Figure 2A, Figure S1A), we compared modulation in the same voxels during neglect and saline sessions. The bar plot shows the attention-related modulation in the ipsilateral (**C**) and contralateral (**D**) ROIs measured as d’ during neglect (SC inactivation) and saline sessions. Error bars indicate 95% CI. Again, we found unilateral effects in aFST/IPa, TPO and FEF ROIs consistent with the comparison between control and during neglect. Because the ROIs were independently identified based on the attention-related modulations during control sessions, these results also address the concern that we might have been biased to see reductions in modulation when comparing control and neglect sessions in the ROIs identified from the control sessions.

**Figure S5. Comparison of the effects of neglect on the attention-related modulations in different cortical ROIs (related to Figures 5 and 6).**

**(A)** Attention-related modulation during midbrain-induced neglect sessions is plotted against attention-related modulation during control sessions for each significant ROI in the ipsilateral hemisphere of monkey #1 for the second-order orientation pulse task. The largest reduction was observed in region aFST/IPa (red arrow).

**(B)** Attention-related modulation for each significant ROI in the ipsilateral hemisphere of monkey #2 during neglect induced by reversible inactivation of the FEF, during the motion-change detection task. Again the largest reduction was observed in region aFST/IPa (red arrow).

