## Supplemental for "Spatial neglect is causally linked to an attention-related area in macaque temporal cortex"

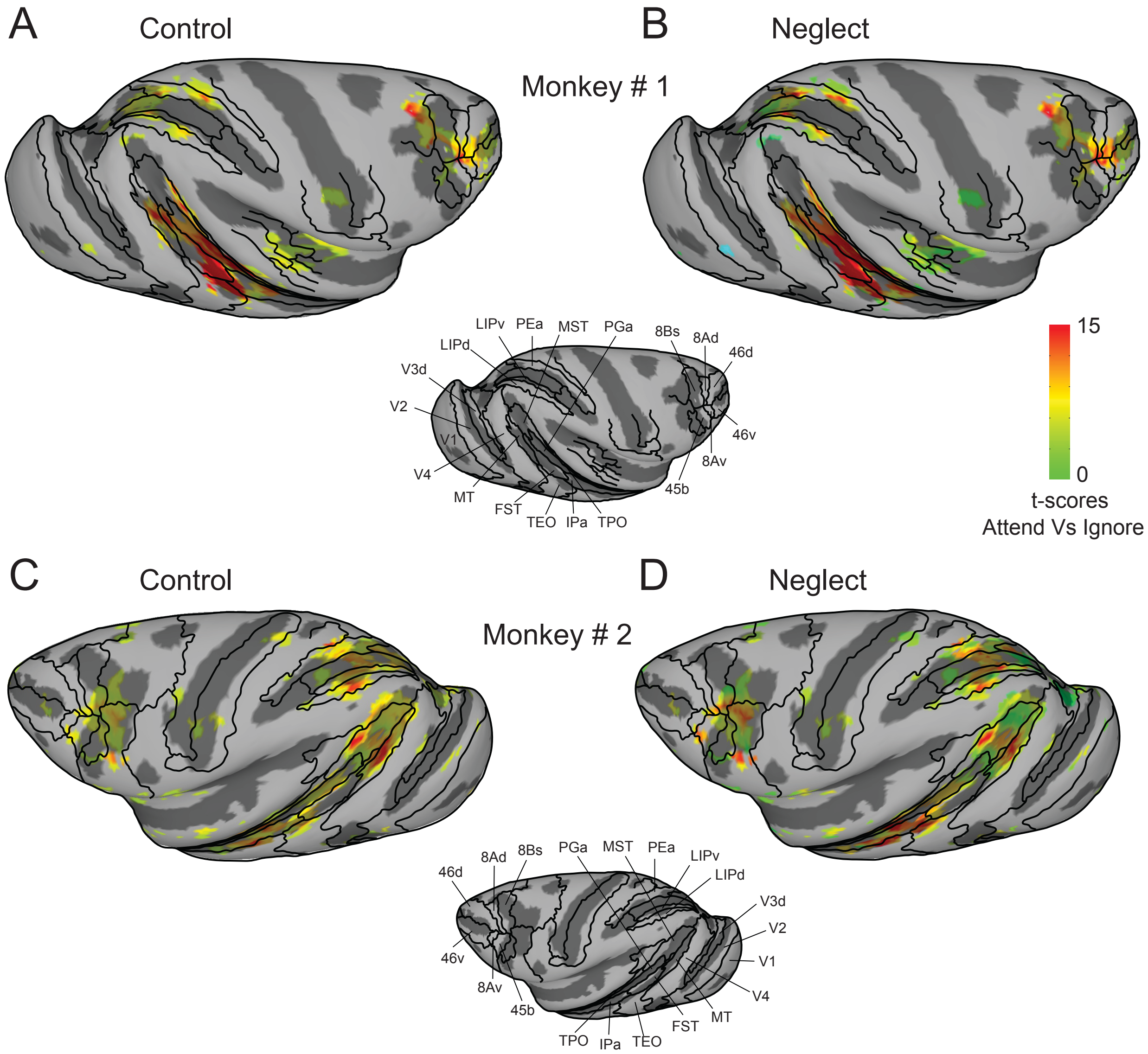


**Figure S1. Cortical maps of attention-related modulation in the contralateral hemisphere during control and neglect sessions (related to Figure 2)**

(**A,B**) Inflated cortical maps of t-scores showing attention-related modulation in contralateral (right) hemisphere of monkey # 1 during control (**A**) and neglect sessions (**B**). The maps were thresholded during control sessions (**A**) and the modulation for the same voxels is shown during the neglect sessions (**B**).

(**C,D**) Same conventions as (**A,B**) but for monkey # 2.

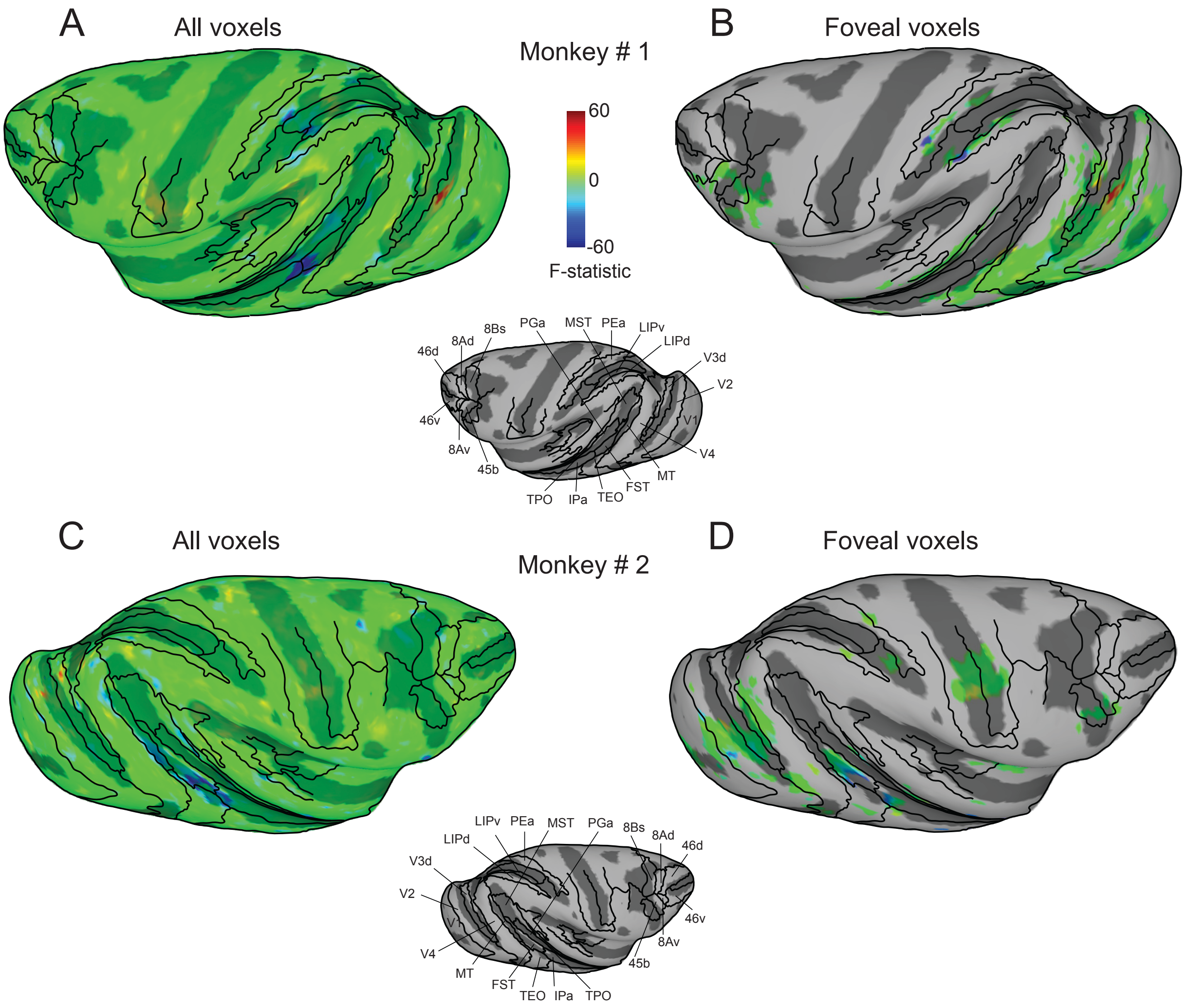


**Figure S2. Effect of midbrain-induced neglect (SC inactivation) on all voxels and voxels that map onto foveal locations (related to Figure 3)**

Inflated cortical maps of interaction term F-statistic show voxels whose modulation was affected during neglect induced by SC inactivation in the ipsilateral hemispheres of both monkeys. The blue color indicates voxels whose modulation was reduced the most during neglect and the green colors indicate voxels with no reduction in modulation. (**A,C**) The effect during neglect is shown for all voxels in the ipsilateral hemisphere of monkey # 1 (**A**) and monkey # 2 (**C**). (**B,D**) The effect during neglect is shown for voxels that map onto foveal locations in the ipsilateral hemisphere of monkey # 1 (**B**) and monkey # 2 (**D**). Cortical regions corresponding to the fovea were unaffected by midbrain-induced neglect following SC inactivation; the main effect of SC inactivation was diminished attention-related enhancement in *non-foveal* regions.

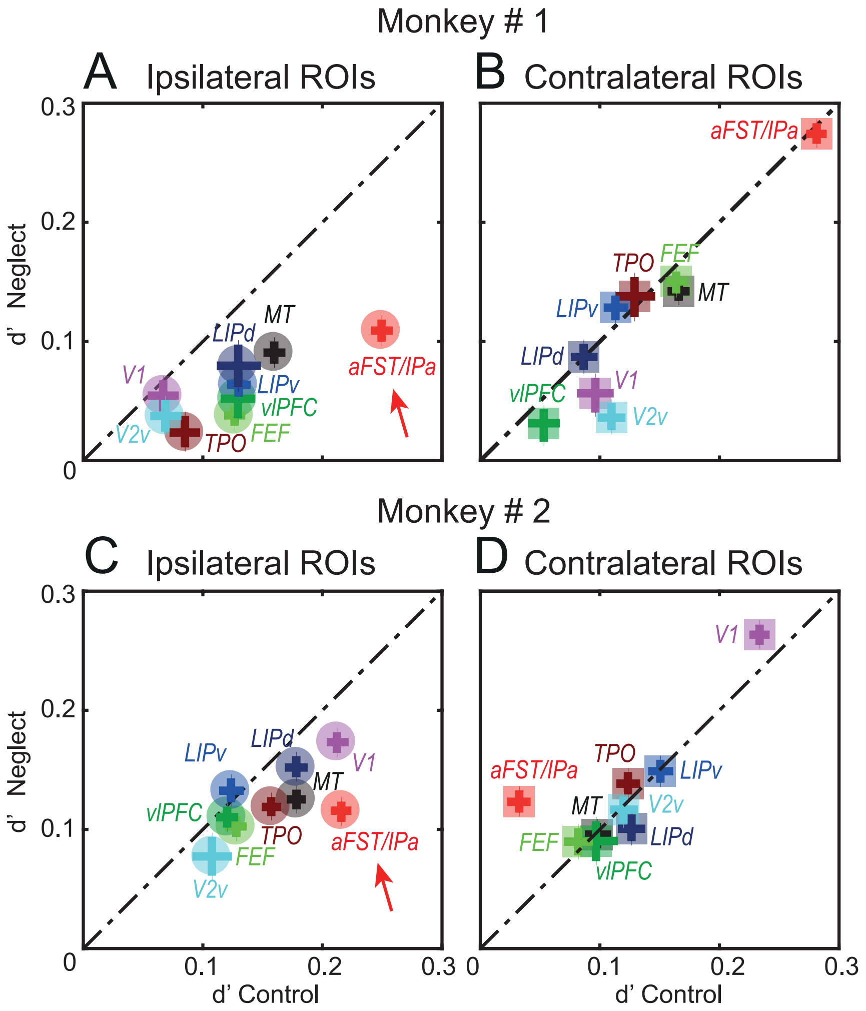


**Figure S3. Comparison of the effects of midbrain-induced neglect on the attention-related modulations in cortical ROIs (related to Figure 4).**

For both monkeys, SC inactivation had the strongest effects on the ipsilateral aFST/IPa region (red arrows). Attention-related modulation during neglect sessions is plotted against the attention-related modulation during control sessions for each ROI in the ipsilateral (**A,C**) and contralateral (**B,D**) hemispheres of both monkeys. In ROIs that lie significantly below the line of identity, attention-related modulation during neglect was significantly reduced compared to the modulation during control. The error bars indicate boot-strapped 95% confidence intervals. Among all ROIs in the ipsilateral hemispheres of monkey #1 (**A**) and monkey #2 (**C**), the aFST/IPa ROI (red symbol and arrows) showed the strongest reduction in modulation and hence lies the furthest from the line of unity slope. In contrast, the aFST/IPa regions in the contralateral hemisphere of monkey #1 (**B**) and monkey #2 (**D**) did not show significant reductions.

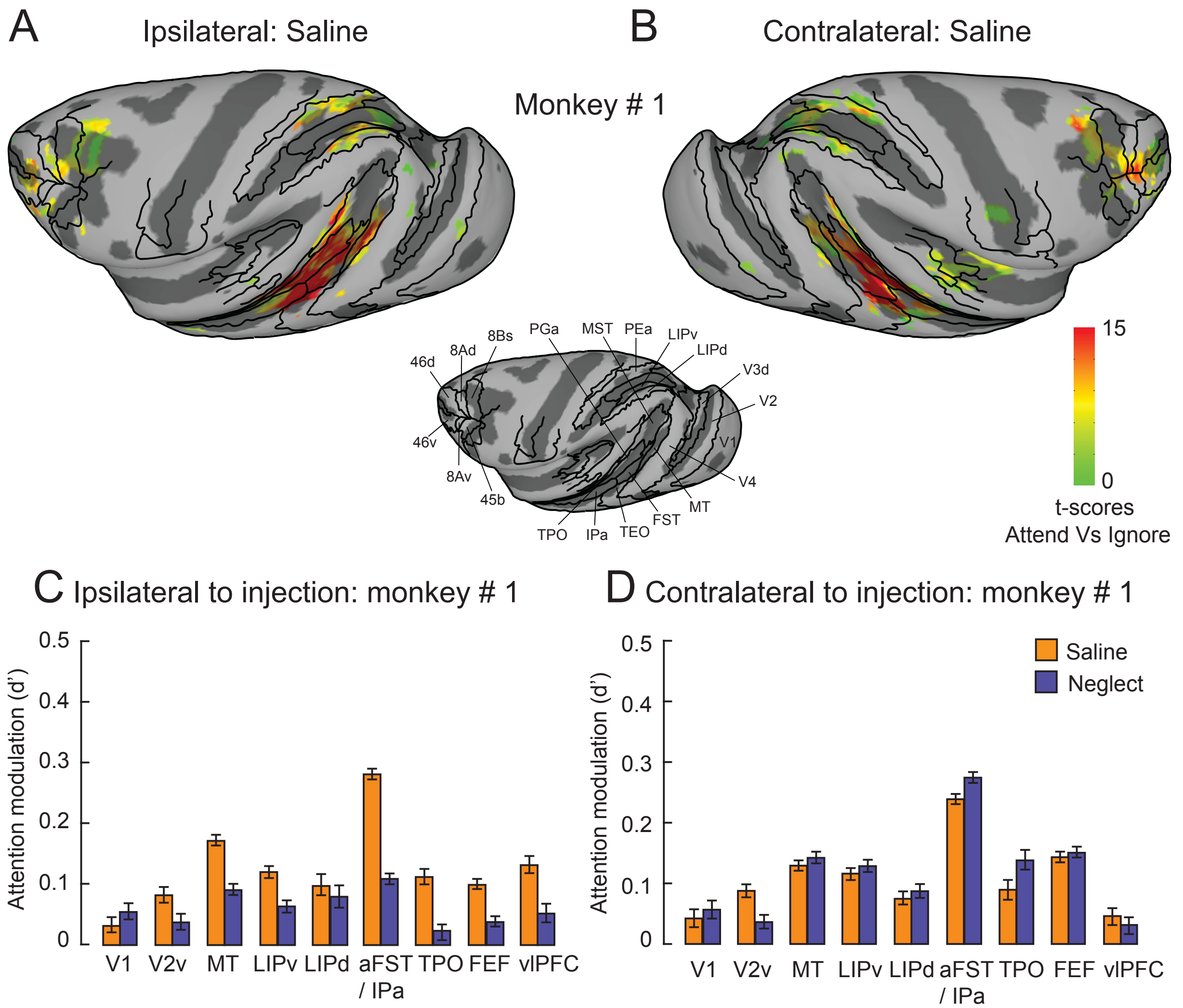


**Figure S4. Comparison of attention-related modulation in cortical ROIs between neglect (SC inactivation) and saline sessions (related to Figure 4)**

(**A,B**) Inflated cortical maps of t-scores showing attention-related modulation in ipsilateral (**A**) and contralateral (**B**) hemispheres of monkey # 1 during saline injection sessions. The maps were thresholded during control sessions (Figure 2, Figure S1) and the modulation for the same voxels is shown during the saline sessions (**A,B**).

(**C,D**) In the ipsilateral and contralateral ROIs defined based on the attention-related modulation map during control (Figure 2A, Figure S1A), we compared modulation in the same voxels during neglect and saline sessions. The bar plot shows the attention-related modulation in the ipsilateral (**C**) and contralateral (**D**) ROIs measured as d' during neglect (SC inactivation) and saline sessions. Error bars indicate 95% CI.

Again, we found unilateral effects in aFST/IPa, TPO and FEF ROIs consistent with the comparison between control and during neglect. Because the ROIs were independently identified based on the attention-related modulations during control sessions, these results also address the concern that we might have been biased to see reductions in modulation when comparing control and neglect sessions in the ROIs identified from the control sessions.

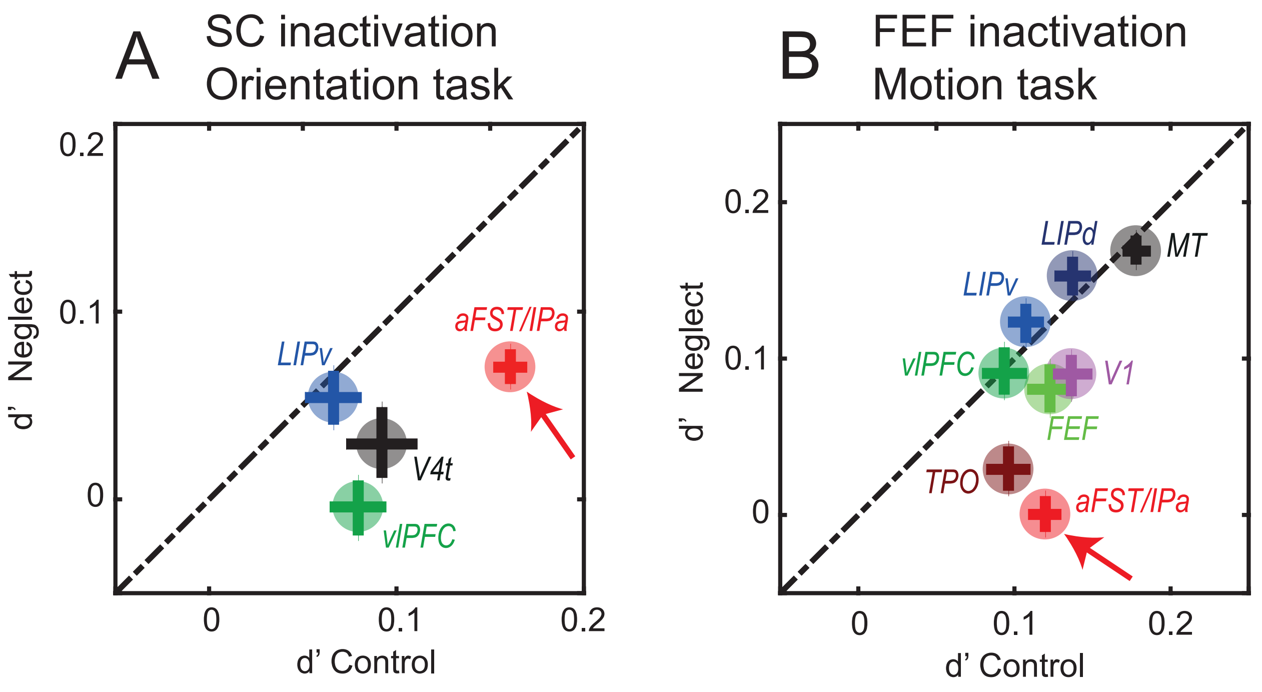


**Figure S5. Comparison of the effects of neglect on the attention-related modulations in different cortical ROIs (related to Figures 5 and 6).**

(**A**) Attention-related modulation during midbrain-induced neglect sessions is plotted against attention-related modulation during control sessions for each significant ROI in the ipsilateral hemisphere of monkey #1 for the second-order orientation pulse task. The largest reduction was observed in region aFST/IPa (red arrow).

(**B**) Attention-related modulation for each significant ROI in the ipsilateral hemisphere of monkey #2 during neglect induced by reversible inactivation of the FEF, during the motion-change detection task. Again the largest reduction was observed in region aFST/IPa (red arrow).
